# Healing potential of wild type and recombinant S100A8 to attenuate skin wound inflammation

**DOI:** 10.1101/2020.09.16.298380

**Authors:** Akinshipo Abdul Warith Olaitan, Lin Chen, Luisa A. DiPietro

## Abstract

**Background:** Wounds represent a major health burden in our society and poorly healing wounds are a significant clinical problem worldwide, During the acute inflammatory, neutrophils which are normal wound scavengers seems to create additional tissue destruction and promote scar formation. This project examined the utility of using pluronic gel to deliver ala^42^S100A8, a peptide that repels neutrophils, to wounds, allowing more regenerative repair.

**Method:** Excisional wound models on female BALB/c mice were made and 4 treatments including pluronic gel only group, Wild type S100A8 (1, 2, and 4μg) with Pluronic gel, and ala^42^S100A8 (1, 2, and 4μg) with Pluronic gel were applied to the wounds. Wounds were harvested at day 1 and day 3. Myeloperoxidase (MPO) protein level was examined using an ELISA kit and cytokine protein expression of CXCL1 (GRO-1), CXCL2 (MIP-2). IL-6, and TNF-α was determined using a multiplex ELISA kit.

**Results:** MPO level in Pluronic gel treated wounds at day 1 was significantly higher than that in control, suggesting that the Pluronic gel itself causes increased inflammation in wounds, while treatment with 1μg of s100A8 or 1 and 4μg of ala42S100A8 seemed to decrease MPO at day 1 compared to the Pluronic gel treated wounds. Treatment with 1μg of s100A8 also led to a decrease IL-6 and TNF-α production at day 1 when compared to the Pluronic gel group, although no statistical difference was observed

**Conclusions:** Our findings strongly suggest that wound inflammation is reduced by treatment with 1ug of S100A8. As such, this study provides proof-of-principal for further investigations of S100A8/ala^42^S100A8 as a wound therapeutic. Additional studies with lower doses and increased sample size, along with the use of alternative delivery systems, will provide important information about the utility of this approach.

## Introduction

Wound healing is a complex yet well-regulated process in which multiple resident cells, recruited inflammatory cells, and stem cells interact to create an environment that supports the repair process. An optimal inflammatory response is a normal and important part of healing that helps to eliminate contaminating microorganisms and dead or injured cells (Eming, Krieg, & Davidson, 2007). Wounds represent a major health burden in our society and poorly healing wounds are a significant clinical problem worldwide (Sen, 2019). Therefore, novel strategies for chronic wound treatment are required. In adult tissues, injury elicits an acute inflammatory response and the role of neutrophils in the acute phase of healing wounds has been demonstrated. Neutrophils, while capable of decontaminating wounds, can cause further tissue destruction primarily via the generation of free oxygen radicals and prolong repair. Moreover, diminished neutrophil activity is a feature of wounds that heal with minimal scar formation. Previous work has shown that neutrophil depletion results in an accelerated wound closure (Dovi, He, & DiPietro, 2003) Therefore reduced neutrophil recruitment and activation might be beneficial for wound closure rates possibly by reducing oxygen demand, wound hypoxia, and bystander damage.

As the level of neutrophils decreases at day 2–4 post-wounding, macrophages become the dominant inflammatory cells in the wound, persisting for approximately 14 days post-wounding (Lucas et al., 2010; Ridiandries, Tan, & Bursill, 2018). Macrophages are key to effective wound healing, as studies from as early as the 1970s have shown that wound healing is significantly delayed if macrophages are depleted (Krzyszczyk, Schloss, Palmer, & Berthiaume, 2018; Leibovich & Ross, 1975). The influx of macrophages to the wound site is stimulated by chemokines such as CCL2, CCL3, and CCL5 (Ridiandries et al., 2018). Macrophages respond to neutrophil signals and help to remove dead and senescent neutrophils through a process called efferocytosis (Koh & DiPietro, 2011). In diabetic wounds, a failure of the macrophages to effectively remove dead neutrophils has been associated with the persistence of neutrophils in the wound and healing impairment (Khanna et al., 2010).

Given the importance of appropriate inflammation to wound healing, many therapeutic approaches to modulate inflammation in order to improve wound repair outcomes have been proposed, and more are under development. One emerging approach is the use of bioengineering constructs to modify the inflammatory response at sites of injury and to facilitate tissue regeneration when tissue is damaged or lost due to trauma, infection, malignancy, surgery, or physical abnormalities. As one example, traditional gel scaffolds such as Poloxamer 407 (Pluronic F-127) can be loaded with molecules that inhibit detrimental inflammation, such as excessive neutrophil activity. Pluronic F-127 is a thermoreversible hydrogel widely used in pharmaceuticals as a carrier for various routes of administration including oral, topical, intranasal, vaginal, rectal, ocular, and parenteral. Pluronic F-127 has a good solubilizing capacity, low toxicity and is, therefore considered a good medium for drug delivery systems. Pluronic F-127 has been widely used in wound healing, mainly as a delivery vehicle for a variety of soluble mediators, including antibodies, cytokines, and growth factors (Mori, Shaw, & Martin, 2008; Dibiase and Rhodes 1996; Mori et al. 2008; Veyries et al. 2000). Pluronic F-127’s ability to gel at higher temperatures allows it to transform from an applied cold solution to solid gel form when heated by body temperature. Upon its application onto skin or injection into a body cavity, the gel acts as a sustained release depot. (Escobar-Chavez et al. 2006).

S100A8 (calgranulin A) and its analogs represent possible anti-neutrophilic molecules which might be effective wound therapeutics. S100A8 (calgranulin A) is an oxidation-sensitive anti-inflammatory protein which combines with S100A9 to form the heterocomplex calprotectin. Calprotectin represents 40% of total protein in the neutrophil cytosolic fraction by weight, and provides important functionality for neutrophils. S100A8 (as well as a related protein, S100A9) inhibits neutrophil chemotaxis toward bacterial products (i.e. formylated peptides) in-vitro through the process of fugetaxis (Sroussi, Berline, Dazin, Green, & Palefsky, 2006; Sroussi, Berline, & Palefsky, 2007). Ala^42^S100A8, an oxidation resistant analog of S100A8 that was engineered using site-directed mutagenesis, retains its chemorepulsive activity under oxidative conditions which would otherwise inactivate the wild type S100A8 protein. In the rat air-pouch model of acute inflammation, ala^42^S100A8 inhibited the recruitment of neutrophils stimulated by bacterial endotoxins (Sroussi et al., 2006).

The goal of the current study was to investigate the effect of wild type and ala^42^S100A8 treatment of acute wounds. We hypothesized that ala^42^S100A8 peptide, delivered using Pluronic hydrogel, would effectively modulate neutrophil function at sites of injury.

## Materials and methods

### Animals

Nine to ten-week old female BALB/c mice were purchased from the Jackson Laboratory (Bar Harbor, Maine). The animals were adapted under controlled temperature (22-24°C) and illumination (12 h light/dark cycles) 1 week before and during the experiments. They had free access to food and water and were kept in individual cages of clean metal. The animals were monitored daily by the animal-care staff and investigators. Any mice displaying signs of infection, severe skin rashes, wasting, and hunching were euthanized. All other mice were sacrificed by lethal CO2 overdose followed by cervical dislocation at the end of each stage of the experiment. All animals received humane care in accordance with the National Institutes of Health’s Guide for the Care and Use of Laboratory Animals (NIH Publication No. 85-23, revised 1996). This protocol was approved by the Committee on Animal Research at the University of Illinois at Chicago.

### Preparation of hydrogels

Hydrogel formulations were prepared as described by Schmolka (Schmolka, 1972). Briefly, 30mg Pluronic F-127 powder (Sigma–Aldrich St. Louis, MO, USA) was slowly added to 120μl chilled phosphate buffer solution (PBS) buffer solution to produce a 25% concentration, then stored in refrigerator (4-5°). This gel mixture was prepared at the time of the experiment and was kept at (4-5°) to maintain liquid form prior to application. Test formulations were prepared by adding a weight percentage of the s100A8 or ala^42^s100A8 to hydrogel while held on ice.

### Excisional wounds

Animals were taken to the surgical room, anaesthetized by an intraperitoneal injection of ketamine (100 mg/kg) and xylazine (5 mg/kg). Subsequently their backs were shaved and cleansed with 70% ethanol. Six 3mm full thickness excisional wounds were made on the shaved dorsal skin on opposite sides of the midline using a 3-mm punch biopsy (Acuderm, Inc., Ft. Lauderdale, FL) and treated as described below. Wounds and surrounding tissues were harvested using 5-mm biopsy punches for samples at day 1 and day 3.

### Wound treatment

The gel application was prepared prior to surgery. The experimental design included 4 groups consisting of control wounds receiving no treatment, Pluronic gel only group, S100A8 (1, 2, and 4μg) with Pluronic gel, and ala^42^S100A8 (1, 2, and 4μg) with Pluronic gel. Wild type S100A8 and ala^42^S100A8 were kindly provided by Dr. Hervé Sroussi (Harvard School of Dental Medicine). The doses of s100A8 and ala^42^s100A8 were selected based upon previous studies (H. Y. Sroussi et al., 2006) S100A8 or ala^42^S100A8 containing hydrogel, or control gel were applied topically to wounds in 10μl of gel solution per wound using a micropipette. Each hydrogel was applied in a fashion that established a uniform thickness covering the wound and extending 1-2mm beyond the edges. Once placed on the wound, the gel solidifies.

### ELISA and multiplex ELISA

Mice were sacrificed on day 1 and day 3 post wounding. Skin wounds harvested, snap frozen, and stored at −80°C until use. Skin wound tissue was homogenized in ice cold PBS with a protease inhibitor cocktail designed for mammalian cells (Protease Inhibitor Cocktail, Catalog number P8340, Sigma Aldrich, St. Louis MO) followed by sonication. Samples were centrifuged at 16,000rpm for 15 minutes. The supernatants were collected and stored at −80°C until use. Myeloperoxidase (MPO) protein level was examined using an ELISA kit (eBioscience, San Diego, CA) and cytokine protein expression of CXCL1 (GRO-1), CXCL2 (MIP-2). IL-6, and TNF-α was determined using a multiplex ELISA kit (eBioscience, San Diego, CA). Assays were performed according to the manufacturers’ instructions.

### Statistical analyses

Data are expressed as mean ± SEM. Multiple t test was used for statistical analysis using GraphPad Prism 5 (GraphPad Software, Inc., La Jolla, CA). n=3 for each treatment group at each time point. P-values less than or equal to 0.05 were considered statistically significant.

## Results

### Neutrophil Content

We first examined whether treatment of wounds with Pluronic F-127 gel +/- S100A8 had an effect on neutrophil content. Myeloperoxidase (MPO) activity was measured as an indicator for neutrophil infiltration. As demonstrated Fig. 1, MPO level in Pluronic gel treated wounds at day 1 was significantly higher than that in control, suggesting that the Pluronic gel itself causes increased inflammation in wounds. Treatment with 1μg of s100A8 or 1 and 4μg of ala^42^S100A8 seemed to decrease MPO at day 1 compared to the Pluronic gel treated wounds (Fig. 1). However, the differences did not reach statistical significance.

**Figure 1:**
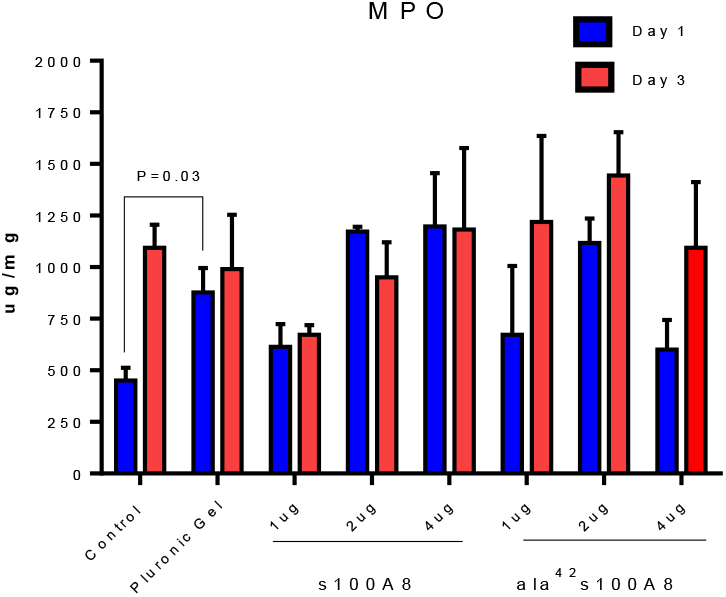
The effect of treatments on neutrophil content. MPO level (ug/ml) is shown for control and treated wounds. P values are shown for significant differences as assessed by a Multiple t test. Bars and lines = means +/ SEM.

### Chemokine Levels: CXCL-1 and CXCL2

Similar to MPO, the levels of neutrophil chemoattractants CXCL1 and CXCL2 in Pluronic gel treated wounds at day 1 and/or day 3 were significantly higher than those in the control untreated group. s100A8 treatment at a dose of 1ug or 2 ug caused an apparent although not significant decrease in CXCL1 at days 1 & 3 and day 1 only, respectively. In general, higher doses of s100A8, and all doses of ala^42^S100A8 seemed to lead to elevated CXCL1 and CXCL2 levels at day 3 (Fig. 2 and 3).

**Figure 2.**
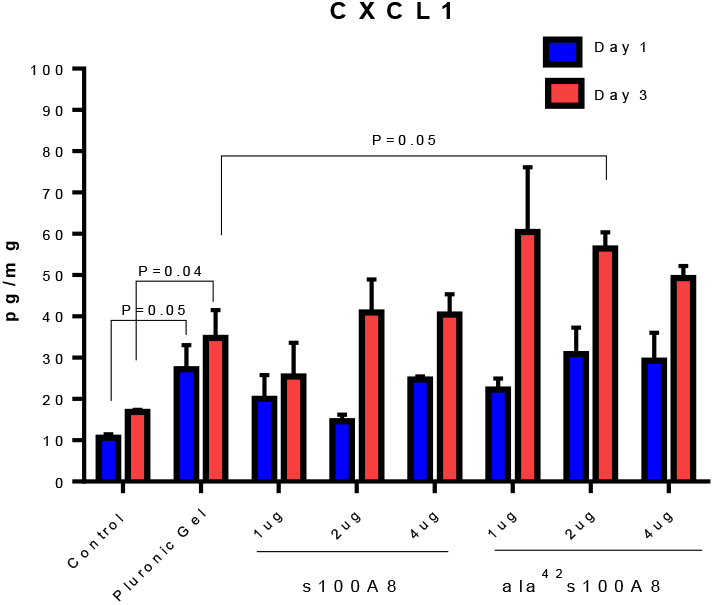
CXCL1 level (ug/ml) is shown for control and treated wounds. P-values show significant differences as assessed by ANOVA and Bonferroni post-hoc test. Bars and lines = means +/ SEM.

**Figure 3.**
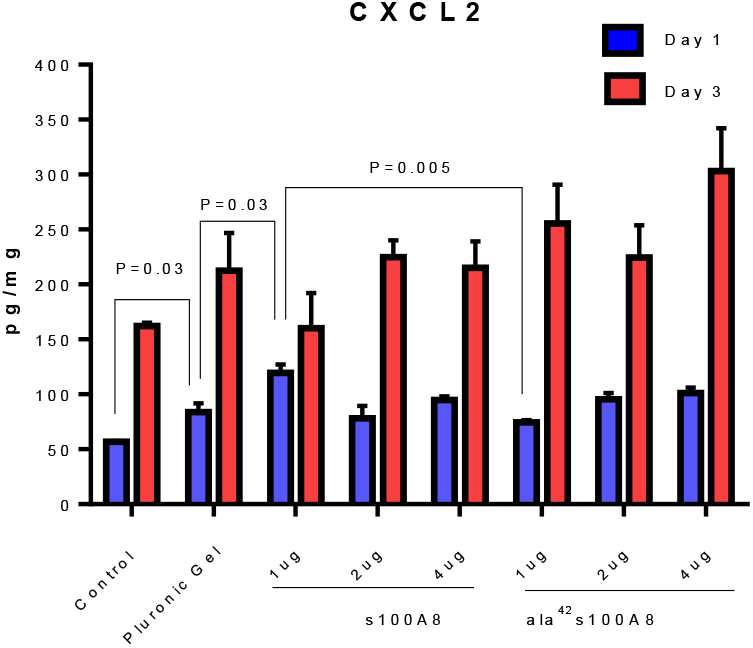
CXCL2 level (ug/ml) is shown for control and treated wounds. P-values show significant differences as assessed by ANOVA and Bonferroni post-hoc test. Bars and lines = means +/ SEM

### IL-6 and TNF-α

Levels of IL-6 and TNF-α were higher in the wounds treated with Pluronic gel than untreated control wounds at day 3. Treatment with 1μg of s100A8 appeared to decrease IL-6 and TNF-α production at day 1 when compared to the Pluronic gel group, although no statistical difference was observed (Fig. 4 and 5).

**Figure 4.**
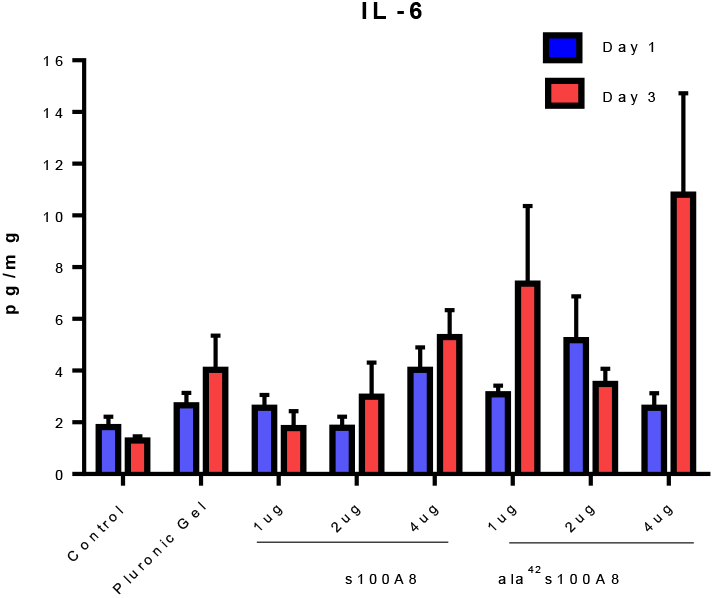
IL-6 level (ug/ml) is shown for control and treated wounds.

**Figure 5.**
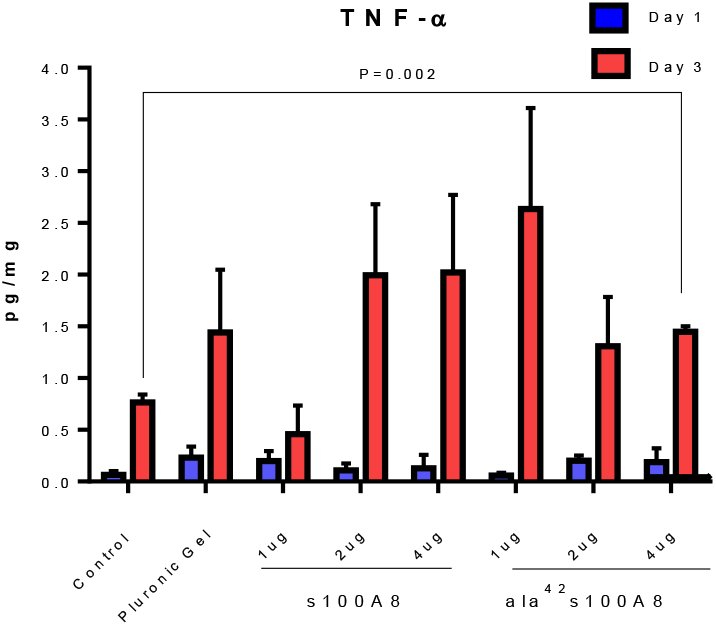
TNF-α level is shown for control and treated wounds. P-values show significant differences as assessed by ANOVA and Bonferroni post-hoc test. Bars and lines = means +/ SEM.

## Discussion

The long-term goal of this work is to develop therapeutic agents that will regulate inflammation, reduce scar formation, and promote ideal tissue regeneration. In the current study, the effect of treatment of wounds with S100A8 or recombinant ala^42^S100A8 protein on neutrophil content, chemokines CCL1 and CCL2, and inflammatory cytokine levels was modest. Treatment of wounds with 1μg of S100A8 or 1 μg and 4μg of ala^42^S100A8 at day 1 led to a modest although not statistically significant decrease neutrophil content.

One important finding of our study is that Pluronic gel alone seems to cause an increase in wound inflammation. The current study demonstrates that the application of Pluronic gel itself can stimulate neutrophilic infiltration in wounds perhaps by way of an increase in CXCL1 and CXCL2 or other cytokines. GRO1/CXCL1 is expressed by macrophages, neutrophils and epithelial cells (Becker, Quay, Koren, & Haskill, 1994) and has neutrophil chemoattractant activity (Schumacher, Clark-Lewis, Baggiolini, & Moser, 1992). Similarly, MIP-2/CXCL2 is also a powerful neutrophil chemoattractant and is involved in many immune responses including wound healing, cancer metastasis, and angiogenesis (Ridiandries et al., 2018). Pluronic gel treated wounds also showed a relative increase in IL-6 and TNF-α, both proinflammatory cytokines with potent effects on healing outcomes. In particular, high TNF-α levels have been linked to poor healing. Although a few studies suggest that TNF-α treatment might improve healing (Frank et al, 2003), most studies show that high levels of TNF-α are detrimental to repair (Ashcroft et al., 2012; Barrientos, Brem, Stojadinovic, & Tomic-Canic, 2014; Mirza, DiPietro, & Koh, 2009). Following injury, TNF-α. is quickly released by endothelial cells, keratinocytes and fibroblasts, initiating the inflammatory phase of healing by promoting inflammatory leukocyte recruitment (Mast & Schultz, 1996). In wounds, a continued cellular source of TNF-α is recruited by neutrophils and macrophages, and this process yields a positive amplification circuit for extending the inflammatory responses (Barrientos et al., 2014). In contrast to TNF-α, IL-6 is a pro-inflammatory that is secreted by macrophages as well as activated T and B lymphocytes (McFarland-Mancini et al., 2010). Prior studies show that topical treatment of wounds with recombinant murine IL-6 delays the healing process (Gallucci et al., 2001), suggesting that excess IL-6 would be detrimental to repair. High levels of IL-6 have been associated with scar formation, as non-scarring fetal wounds contain decreased IL-6, while keloid scars have high levels of this cytokine (Ghazizadeh, 2007; Zgheib, Xu, & Liechty, 2014). Thus, an increase in IL-6, if induced by Pluronic gel, would be likely to be detrimental to healing outcomes.

The use of recombinant proteins, such as CCL2 (MCP-1), to modify the inflammatory reponse in wounds has been shown to be beneficial in many experimental situations (Öhnstedt et al, 2019; Dipietro et al, 2001) In the current study, the failure of of s100A8 and ala^42^s100A8 treatment to successfully reduce wound inflammation may be due to several reasons. A likely cause seems to be the delivery system, as the Pluronic gel appears to negatively influence wound inflammation in and of itsself. Moreover, Pluronic gel can absorb to the surface of the wound, impairing the diffusion of the recombinanat protein to the wound interior (Kant et al., 2014). While the negative effect of Pluronic gel might be small, the impact could be important to the use of this agent as a delivery vehicle. Setting aside the limitations of the delivery system, our study does suggest that lower doses of S100A8 might be beneficial to repair. We observed that the lowest dose of 1ug of S100A8 reduced inflammation when compared to Pluronic gel, whereas higher doses of S100A8 and alaS100A8 showed variable levels of inflammation when compared to the gel alone. Another caveat to the results is the time frame of investigation. Earlier measurements, such as 6 hours post wounding (a time that corresponds to the initial burst phase of cytokine release), might demonstrate a larger effect of the treatments. Finally, our experiments did not address any effects on macrophage function, a parameter that certainly needs to be investigated in future studies.

In conclusion, our findings strongly suggest that wound inflammation is reduced by treatment with 1ug of S100A8. As such, this study provides proof-of-principal for further investigations of S100A8/ala^42^S100A8 as a wound therapeutic. Additional studies with lower doses and increased sample size, along with the use of alternative delivery systems, will provide important information about the utility of this approach.

## Acknowledgements

This study was supported by grant from NIH/FORGATY INTERNATIONAL D43TW010134 Building research and Innovation in Nigerian Sciences (BRAINS) (to AAWO). We thank Dr. Hervé Sroussi (Harvard School of Dental Medicine) for the kind gift of s100A8 and ala^42^s100A8, and Mr. Junhe Shi (UIC Center for Wound Healing and Tissue Regeneration) for technical support. AAWO appreciates the input of his mentors Prof. WA Adeyemo (University of Lagos) and Prof. Niyi Oshuntoki (University of Lagos).

